# Landscape connectivity alters the evolution of density-dependent dispersal during pushed range expansions

**DOI:** 10.1101/2021.03.03.433752

**Authors:** Maxime Dahirel, Aline Bertin, Vincent Calcagno, Camille Duraj, Simon Fellous, Géraldine Groussier, Eric Lombaert, Ludovic Mailleret, Anaël Marchand, Elodie Vercken

## Abstract

As human influence reshapes communities worldwide, many species expand or shift their ranges as a result, with extensive consequences across levels of biological organization. Range expansions can be ranked on a continuum going from pulled dynamics, in which low-density edge populations provide the “fuel” for the advance, to pushed dynamics in which high-density rear populations “push” the expansion forward. While theory suggests that evolution during range expansions could lead pushed expansions to become pulled with time, empirical comparisons of phenotypic divergence in pushed vs. pulled contexts are lacking. In a previous experiment using *Trichogramma brassicae* wasps as a model, we showed that expansions were more pushed when connectivity was lower. Here we used descendants from these experimental landscapes to look at how the range expansion process and connectivity interact to shape phenotypic evolution. Interestingly, we found no clear and consistent phenotypic shifts, whether along expansion gradients or between reference and low connectivity replicates, when we focused on low-density trait expression. However, we found evidence of changes in density-dependence, in particular regarding dispersal: populations went from positive to negative density-dependent dispersal at the expansion edge, but only when connectivity was high. As positive density-dependent dispersal leads to pushed expansions, our results confirm predictions that evolution during range expansions may lead pushed expansions to become pulled, but add nuance by showing landscape conditions may slow down or cancel this process. This shows we need to jointly consider evolution and landscape context to accurately predict range expansion dynamics and their consequences.

## Introduction

The distribution ranges for many species are currently shrinking, shifting or expanding as a direct or indirect result of human influence. Describing, understanding and predicting these changes and their consequences is currently the focus of substantial research effort, in particular in climate-tracking or invasive species (Chuang & Peterson, 2016; Lenoir et al., 2020; Renault et al., 2018). Given the key role of within-species trait variability for ecosystem functioning (Des Roches et al., 2018; Jacob et al., 2019; Little et al., 2019; Raffard et al., 2019; Violle et al., 2012), knowing how phenotypes are redistributed in space during range expansions and range shifts is important to understand the ecological and evolutionary dynamics at play in the resulting communities (Cote et al., 2017; Miller et al., 2020; Renault et al., 2018).

The speed at which a species’ range expands in space is, ultimately, a function of both population growth and dispersal (Lewis et al., 2016). As populations/species differ qualitatively in their growth and dispersal functions (Fronhofer et al., 2018; Gregory et al., 2010; Harman et al., 2020; Sibly & Hone, 2002), due to intrinsic and/or environmental drivers, we can expect them to differ in the way they advance during range expansions too. Building on the framework of reaction-diffusion equations, one can discriminate between “pushed” and “pulled” expansions (Lewis et al., 2016; Stokes, 1976), although it may be more accurate to think of it as a continuum of “pushiness” (Birzu et al., 2018). Pulled expansions are the type often implied “by default” in many ecological studies (Deforet et al., 2019; see e.g. Weiss-Lehman et al., 2017). Pulled expansions assume dispersal and growth are either constant or maximal at the lowest densities. This leads to expansions being “pulled” forward by the few individuals at the low-density, recently populated edge (Lewis et al., 2016; Stokes, 1976). However, in many species, dispersal is actually more likely at high densities, as a way to escape increased competition (Harman et al., 2020; Matthysen, 2005). Additionally, populations can exhibit Allee effects (Allee & Bowen, 1932; Courchamp et al., 2008), i.e. have their growth rate decrease at lower densities. In both cases, this leads to the product of per capita growth and dispersal being highest at intermediate or high densities; these expansions are thus “pushed” by older populations that have reached these densities, instead of being primarily driven by low-density edge populations.

Individuals founding new populations at the leading edge of an expansion are likely a non-random sample of available phenotypes, because individuals with traits facilitating spread are more likely to reach these new habitats in the first place. If these individual differences are heritable, then these traits can evolve during expansion, as phenotypes facilitating spread accumulate at the expansion edge with time (Cwynar & MacDonald, 1987; Phillips & Perkins, 2019; Shine et al., 2011). Evolution of increased dispersal ability in leading-edge populations is now well documented, both in experimental and natural contexts (Chuang & Peterson, 2016; Deforet et al., 2019; Fronhofer et al., 2017; Weiss-Lehman et al., 2017). In addition, relaxed density-dependence at the lower-density edge can select for faster life-history, e.g. higher fecundity (Burton et al., 2010; Van Petegem et al., 2018). Both models and reshuffling experiments (where individuals’ locations are regularly randomized to stop spatial evolution) have demonstrated how these evolutionary changes can accelerate expansions (Perkins et al., 2013; Schreiber & Beckman, 2020; J. M. Travis & Dytham, 2002; Van Petegem et al., 2018; Weiss-Lehman et al., 2017). However, summarizing empirical studies also shows that these directional shifts in population growth, dispersal or associated traits do not always happen during range expansions (Chuang & Peterson, 2016; Merwin, 2019; Van Petegem et al., 2018; Wolz et al., 2020). We need a better understanding of what determines whether or not this evolution will occur, and whether it will affect growth traits or dispersal traits, if we want to successfully predict (and potentially manage) the ecological and evolutionary dynamics of range expansions or shifts.

The position of an expansion on the pushed-pulled continuum can have consequences on its evolutionary dynamics: for instance, (more) pushed expansions should conserve more genetic diversity (Birzu et al., 2018, 2019; Roques et al., 2012). While this effect of expansion type on neutral evolution has been confirmed experimentally (e.g. Gandhi et al., 2019), the possibility that pushed and pulled expansions may also differ in their adaptive evolutionary dynamics has remained almost completely unstudied so far (Birzu et al., 2019). Exploring this is, in our opinion, the next step in pushed expansion studies, given that the distinction between pushed and pulled expansions rests, at its core, on traits (dispersal and fecundity) we now know can evolve during range expansions. Moreover, there is evidence that evolution during range expansion can lead to changes in not only average dispersal between core and edge populations, but also in the density dependence of dispersal, i.e. precisely one of the characteristics that determine whether an expansion is pushed or not. The few (theoretical and empirical) studies that are available hint that evolution at range edges may lead pushed expansions to become pulled (Erm & Phillips, 2020), as they show an initial positive density-dependence in growth or dispersal is lost during expansion (Erm & Phillips, 2020; Fronhofer et al., 2017; Travis et al., 2009; Weiss-Lehman et al., 2017; but see Mishra et al., 2020).

In the current context of habitat loss and fragmentation, several studies have set to explore how habitat connectivity can affect range expansion speeds and/or the evolution of dispersal and other traits during range expansions (Gralka & Hallatschek, 2019; Hunter et al., 2021; Lutscher & Musgrave, 2017; Pachepsky & Levine, 2011; Urquhart & Williams, 2021; Williams, Snyder, et al., 2016; Williams, Kendall, et al., 2016; Williams & Levine, 2018). For instance, using experimental expansions, Williams et al. (2016) showed that evolution had stronger effects on range expansion speeds in patchier landscapes where connectivity was lower (or, conversely, that evolution dampened the negative effects of low connectivity on speed). Experiments and models show that less connected landscapes also select more strongly for large individuals/more competitive individuals than continuous landscapes during expansions, an indication that evolution at expanding range edges can itself be shaped by landscape connectivity (Williams, Snyder, et al., 2016; Williams, Kendall, et al., 2016). Williams and Levine (2018) showed that the effects of density-dependence on expansion speed could be of the same magnitude than those of connectivity, matching theoretical predictions made earlier (Pachepsky & Levine, 2011). However, this study used negative density-dependent dispersal, and as such we cannot directly transpose its results to the study of pushed expansions. In addition, all these studies either focused on a simple, density-independent dispersal trait or, when they did account for density-dependent dispersal, ignored the effects of evolution. As a result, key questions remain, that are important for our ability to successfully predict expansion dynamics: how does connectivity shape the evolution of density-dependent dispersal during expansions? And do connectivity-induced differences in selection pressures influence the stability of an expansion type (pushed or pulled) through time (Birzu et al., 2019; Erm & Phillips, 2020)?

Here we revisit a previous study of experimental range expansions using *Trichogramma* parasitic wasps as a model (Dahirel et al., 2021), in which we showed that reducing landscape connectivity led to increased “pushiness.” Using this data we examine the phenotypic changes underlying the different types of range expansions, in space and time. We first ask whether body size, a trait that is linked to fitness in *Trichogramma* (Durocher-Granger et al., 2011), differs between core and edge populations and across connectivity treatments. We then conduct a common-garden experiment, using the descendants of the expansion experiments, to study whether different range expansion contexts led to contrasted evolutionary changes in traits directly linked to spread, namely dispersal, activity and reproductive success, with special attention to changes in density-dependence in part of the experiments.

## Methods

### Study species and range expansion experiment

This experimental protocol for the expansions is described in detail in a previous article (Dahirel et al., 2021); we here summarise its most relevant aspects.

*Trichogramma* wasps are small (body length ≈ 0.5 mm when adult) egg parasitoids that are relatively easy to maintain on standardised resources in the lab. We used three laboratory “strains” of *Trichogramma brassicae* Bezdenko, 1968 (Hymenoptera: Trichogrammatidae) for our experiment (**Fig. 1A**). Each strain was obtained by mixing three pre-existing isoline populations using Fellous et al. (2014)’s protocol to ensure similar genetic representation of the isolines in the final mixes. Isolines were themselves derived from individuals collected in different sites across western Europe in 2013. The three resulting mixed strains had broadly similar levels of genetic diversity at the start of the experiment, with expected heterozygosity based on 19 microsatellite loci in the 0.3-0.4 range (Dahirel et al., 2021). They were raised using irradiated eggs of the Mediterranean flour moth *Ephestia kuehniella* Zeller 1879 (Lepidoptera: Pyralidae) as a substitution host (St-Onge et al., 2014).

**Figure 1.**
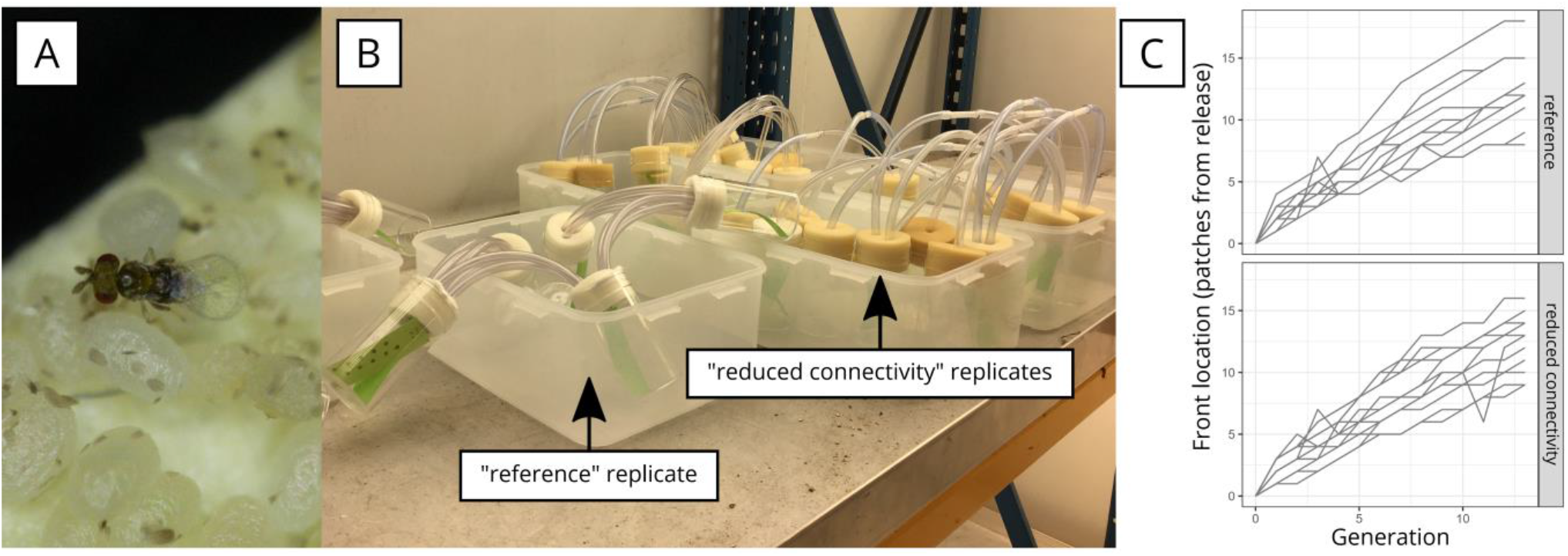
A: *Trichogramma brassicae* on *Ephestia kuehniella* eggs (picture by Géraldine Groussier). B: replicate landscapes used in the range expansion experiment. Picture (by Aline Bertin) shows both reference landscapes (patches connected by three 20 cm tubes) and “reduced connectivity” landscapes (patches connected by one 40 cm tube). Clusters of host eggs on paper strips can be seen in patches. C: Front location (i.e. farthest populated patch) through time for each replicate landscape (data from Dahirel et al., 2021).

We monitored *T. brassicae* spread in 24 experimental linear landscapes (8 per genetic strain) for 14 non-overlapping generations (Generations 0-13, with initially released adults counted as Generation 0, and the experiment stopped at the emergence of Generation 13 adults). Landscapes were made of plastic vials (10 cm height, 5 cm diameter) connected to their nearest neighbours by flexible tubes (internal diameter 5 mm). In half of the replicate landscapes, patches were connected by three 20 cm long tubes (“reference” connectivity). In the other half, connectivity was reduced and patches were only connected by one longer (40 cm) tube (**Fig. 1B**). Patches contained approximately 450 *Ephestia* eggs, on paper strips to facilitate handling, renewed every generation at adult emergence. We started landscapes by placing ≈ 300 unsexed adult wasps in one extremity patch (expansion was only possible in one direction), a number close to the expected equilibrium population size in such a system (Morel-Journel et al., 2016). Each generation, adult individuals were allowed to disperse, mate and lay eggs for 48 hours before they were removed. The landscapes with reduced connectivity had on average more pushed dynamics than the “reference” ones, drawing on both direct (genetic) and indirect arguments (Dahirel et al., 2021). The average expansion speed was similar between the two connectivity treatments (**Fig.1C**, Dahirel et al., 2021). Experimental landscapes, as well as subsequent experiments described below, were kept under controlled conditions (23°C, 70% relative humidity, 16:8 L:D).

### Phenotypic measurements

For our analysis of trait change, we focused on descendants of individuals born towards the end of the experiment in “core” patches (here, the release patches or their immediate neighbours, x = 0 or 1) or in the corresponding “edge” patches (i.e. the farthest populated patch in a landscape at the time of sampling, or the farthest two if there were not enough individuals in the farthest one). We compared them to wasps from the “stock” populations initially used to start the experimental expanding landscapes. Note that mentions of “Xth generation” wasps below indicate the number of generations of experimental landscape expansion before sampling/ transfer to common garden conditions. For some traits (short-term movement, and fecundity and dispersal during the density-independent tests), data was also collected on one or two intermediate generations. For consistency and simplicity, we only analysed (and described below) the latest tested generation for each trait, but made available all data, including intermediate samples (**Data accessibility**).

#### Wasp size

To determine whether landscape connectivity and expansion had an effect on body size, we selected female wasps from the stock populations, and compared them to 12th generation females from the experimental landscapes. Due to logistical constraints, the latter were selected in 8 edge-core pairs of populations (see **Fig. 2** for how they were distributed among landscape treatments and strains). Adding the three stock populations, and accounting for the fact one edge-core pair was only sampled in the core due to limited numbers in the edge population, we measured 316 (91 to 116 per strain) wasps in 18 populations (mean ± SD: 17.6 ± 4.0 wasps per population).

**Figure 2.**
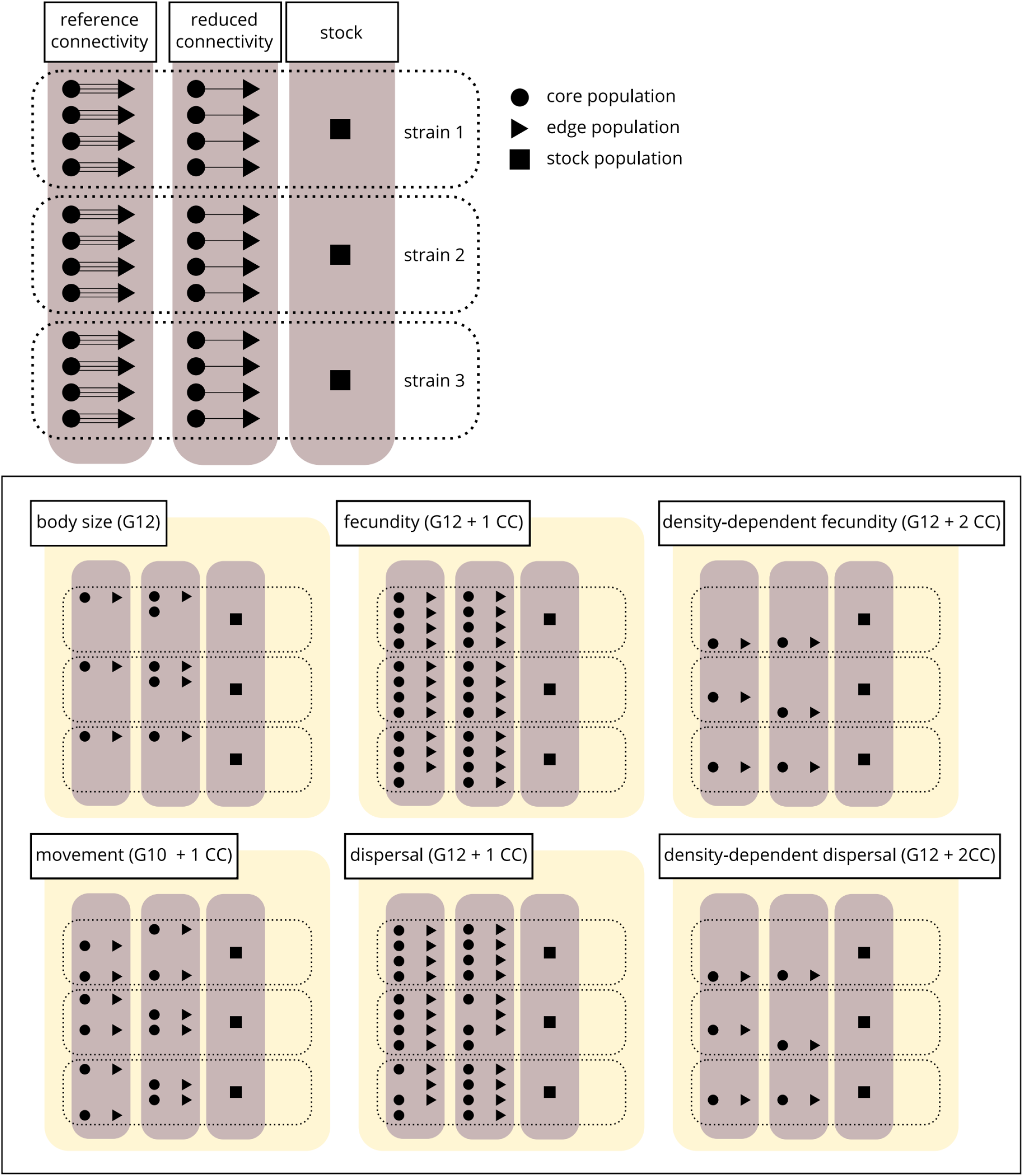
Top: summary of the experimental populations available to sample. Core and edge populations from 24 experimental landscapes (split in two connectivity treatments and three “strains”) were available, along with the corresponding stock populations. Bottom: distribution of the populations actually sampled for each phenotypic trait. The generation at which wasps were taken from the experimental landscapes (+ the number of common garden generation _CC_ before testing) is indicated in parentheses besides the name of each trait.

Wasps were kept in 70% ethanol before phenotypic measurements. We used hind tibia length (in μm) as a body size proxy (e.g. Durocher-Granger et al., 2011). We used a Zeiss AxioImager Z1 microscope equipped with a 40x/0.75 objective to photograph tibias after dissection. Images were managed and measurements done using the OMERO platform (Allan et al., 2012). Wasps were measured by two independent observers; inter-observer agreement was good but not perfect (*r* = 0.93). We thus decided to use a hierarchical approach to explicitly include measurement error in-model (see **Statistical analysis** below) rather than averaging observations before fitting.

#### Short-term movement

To study differences in short-term movement between treatments and between core and edge patches, we analysed F1 offspring of 10th generation wasps, and compared them to each other and to wasps from the stock populations. To control for population density (and other) variations among landscapes, we used a common-garden protocol: wasps removed from their natal landscapes after the egg-laying phase were allowed to lay eggs on new host egg strips for 48h (with ≈ 20 females per ≈ 450 host eggs, i.e. low density conditions). Emerging offspring (unsorted by sex) were placed in an empty and lit 15 × 19 cm rectangular arena, 2 cm high, sealed above and below with a glass sheet. Groups of 15.8 individuals on average (SD: 3.3) were introduced per replicate trial, and their movements filmed for five minutes. To reduce behavioural changes at the edge of the arenas, and their effect on our metrics, we only tracked individuals within a central 7×11 cm area, and the outer parts of the arena were kept in the dark to discourage individuals from approaching the edges. We studied 27 populations (core and edge from 12 of the 24 experimental landscapes + the three stock populations, see **Fig. 2**), with 8 replicate trials per stock population, and 16 replicate trials for each of the remaining populations (except one with only 15), for a total of 119 replicate groups. Video files were analysed using Ctrax (Branson et al., 2009) for tracking and the `trajr` R package (McLean & Volponi, 2018) for computation of movement statistics from trajectories. Most individuals were not tracked continuously for the entire five minutes due to either leaving the filmed area or the loss of individual identity information. As a result, output data were in the form of a series of “tracklets” (i.e. any continuous sub-track longer than 2 seconds), that could not be assigned to a specific individual, only to a specific replicate trial. We therefore first computed metrics at the tracklet level, and then averaged them, weighted by tracklet duration, to generate replicate-level metrics. We used the proportion of total tracked time individuals were active, the average speed and the average sinuosity (Benhamou, 2004). All three movement metrics responded similarly to the experimental protocol; for simplicity, we only present and discuss results from the “proportion of time active” metric here, and models for the other metrics are included in the associated analysis code (see **Data accessibility**).

#### Effective dispersal

F1 offspring of 12th generation wasps (reared in a low-density common-garden setting as described above) were used to evaluate dispersal differences between treatments. We placed groups of 50 unsexed newly emerged wasps in a departure vial connected to an arrival vial by one 40 cm flexible tube (i.e. reduced connectivity conditions). Both vials contained 90 host eggs. We tested 47 populations (core and edge populations from all 24 experimental landscapes, excluding four populations, plus the three stock populations; see **Fig. 2**), with two replicates per “experimental landscape population” and 4 replicates per “stock population,” for a total of 100 replicates (44 × 2 + 3 × 4). One of these replicates was lost, so the final number was 99 replicates. We let wasps in vials for 24h, removed them, then waited 7 days and counted darkened host eggs (an indication of successful parasitoid development). We used the proportion of parasitized eggs found in the arrival patches, relative to the total parasitized eggs in a replicate (departure and arrival patches), as our measure of dispersal rate. As such, it is important to note it is not a measure of the percentage of individuals that dispersed (as dispersers and residents may differ in sex-ratio, fecundity, competitive ability and survival, Ronce & Clobert, 2012), but rather a context-specific measure of effective dispersal or gene flow. This experiment is therefore complementary to the short-term movement experiments (see above), since the former experiment allows us to examine how connectivity and expansion influence individuals’ movement behaviour, and this dispersal experiment allow us to examine their net effect on all three phases of dispersal together (emigration probability, movement/transience, settlement).

#### Fecundity

We placed newly emerged and presumably mated females (obtained at the same time and in the same way as the ones used to measure dispersal) individually in vials containing 90 host eggs, and let them lay eggs for 24h. We then counted the number of darkened host eggs after 7 days as our measure of reproductive success. Because superparasitism (more than one egg per host) frequently happens in *Trichogramma* wasps (Corrigan et al., 1995), this is not a measure of eggs produced stricto sensu, but rather a metric of reproductive success (in most cases, a single adult emerges per host, even when superparasitism occurs; Corrigan et al., 1995). A total of 492 F1 females coming from 50 populations were used, (core and edge from all 24 experimental landscapes _excluding one edge population due to low sample size_ + the three stock populations, see **Fig. 2**) with 9.8 individuals per population on average (SD: 3.3).

#### Density-dependent dispersal and fecundity

F2 descendants of the 12th generation to emerge from experimental landscapes (and from a new set of stock population wasps) were subjected to the same dispersal and reproduction experiments as F1 wasps, with the difference that developmental density conditions before the experiments were manipulated. For these experiments, due to logistic constraints, we studied wasps coming from each of the three stock populations and one randomly selected landscape per connectivity × genetic strain combination (**Fig. 2**). High-density wasps were obtained by placing ≈ 90 F1 parasitized eggs close to maturity with ≈ 90 fresh host eggs; this in effect mimics the conditions in core patches during the expansions, with populations at carrying capacity and a 1 to 1 replacement of host eggs from one generation to the next. Low-density wasps were obtained by placing ≈ 90 parasitized eggs with ≈ 450 fresh hosts; these conditions are closer to the conditions experienced at the range edge. Higher densities likely led to higher superparasitism and higher within-host competition during early development (Corrigan et al., 1995; Durocher-Granger et al., 2011). 341 F2 females were tested in total for the reproductive success experiment (N = 19 or 20 per density level for each of the three stock populations, while 9.3 females were tested on average for the other population × density combinations (SD: 1.7)). For the dispersal experiment, we used 72 groups of 50 wasps, with 4 replicates per stock population × density (4 replicates × 2 densities × 3 strains = 24), and 2 replicates per remaining population × density combination (2 replicates × 2 densities × 2 locations _core/edge_ × 2 connectivity treatments × 3 strains = 48).

### Statistical analyses

Analyses were done using R, versions 4.0.4 and 4.1.0 (R Core Team, 2021). We analysed data using a Bayesian framework using the `brms` R package (Bürkner, 2017) as a frontend for the Stan language (Carpenter et al., 2017). We mostly relied on the `tidybayes` (Kay, 2019), `bayesplot` (Gabry et al., 2019), `patchwork` (Pedersen, 2019) packages, and on the `tidyverse` suite of packages (Wickham et al., 2019), for data preparation, model diagnostics and plotting. We ran four Markov chains per model; the number of iterations per chain was model-dependent (4000 or 8000, with the first half of each chain used as warm-up in either case), and set to be large enough to ensure convergence 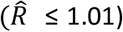 and satisfactory effective sample sizes (both bulk- and tail-effective sample sizes *sensu* Vehtari et al., 2020 > 1000). When posteriors are summarised, all credible/compatibility intervals given are highest posterior density intervals. Priors were chosen to be weakly informative and mostly follow suggestions by McElreath (2020); they are described in detail in **Supplementary Material S1**, along with a formal description of each model.

We used (generalized) linear mixed models to analyse how phenotypic traits (size, short-term movement, reproductive success and effective dispersal) varied between connectivity treatment × location combinations (five levels). We used random effects (random intercepts) of genetic strain, experimental landscape nested in strain, and source location (stock, edge or core patch) nested in landscape to account for phylogenetic relatedness/ shared ancestry among populations (Clutton-Brock & Harvey, 1977; Hadfield & Nakagawa, 2010).

- We used a Gaussian model for size, with tibia length (centred and scaled to unit 1 SD) as the response. In addition to the fixed effect of connectivity × location and the “phylogenetic” random effects described above, and because individuals were measured twice, this model included a random effect of individual identity, allowing us to split (within-population) individual variation from (residual) observation error.
- We analysed the percentage of time active per test group as a function of connectivity × location and phylogeny using a Beta model.
- We analysed reproduction data (number of eggs successfully parasitized) using zero-inflated negative binomial models, as initial analyses revealed zero-inflation. The submodels for the probability of excess zeroes *p* (i.e. reproductive failure) and for the number of eggs otherwise (λ) both included effects of phylogeny and connectivity × location. For simplicity, we do not discuss in the Results section the two submodels separately, but only the overall posterior average fecundities (1 − *p*) × λ. The density-dependent experiment was analysed using a very similar model, with added fixed effects of density and density × connectivity × location interactions.
- Finally, we analysed effective dispersal rates using binomial models. As for fecundity, models included effects of phylogeny and connectivity × location (+ density and density × connectivity × location effects for the density-dependent experiment). Initial models presented some evidence of overdispersion. This was accounted for by adding the total number of eggs laid (centred and scaled to unit 1 SD) as a covariate: while it may indicate a dispersal-fecundity syndrome, a positive link between effective dispersal and total fecundity is also very likely to arise “artificially” in our setup simply because once the departure patch is saturated, individuals can only successfully reproduce if they disperse. Note that in *Trichogramma*, we expect a priori such saturation to appear well below the nominal limit based on host number, due to competition (Dahirel et al., 2021; Morel-Journel et al., 2016). The main conclusions we derive from the model do not change if we do not control for the total number of eggs laid.

## Results

Average tibia length did not differ meaningfully between connectivity treatments and locations (**Fig. 3**, see **Supplementary Figure S.2.1** for pairwise comparisons).

**Figure 3.**
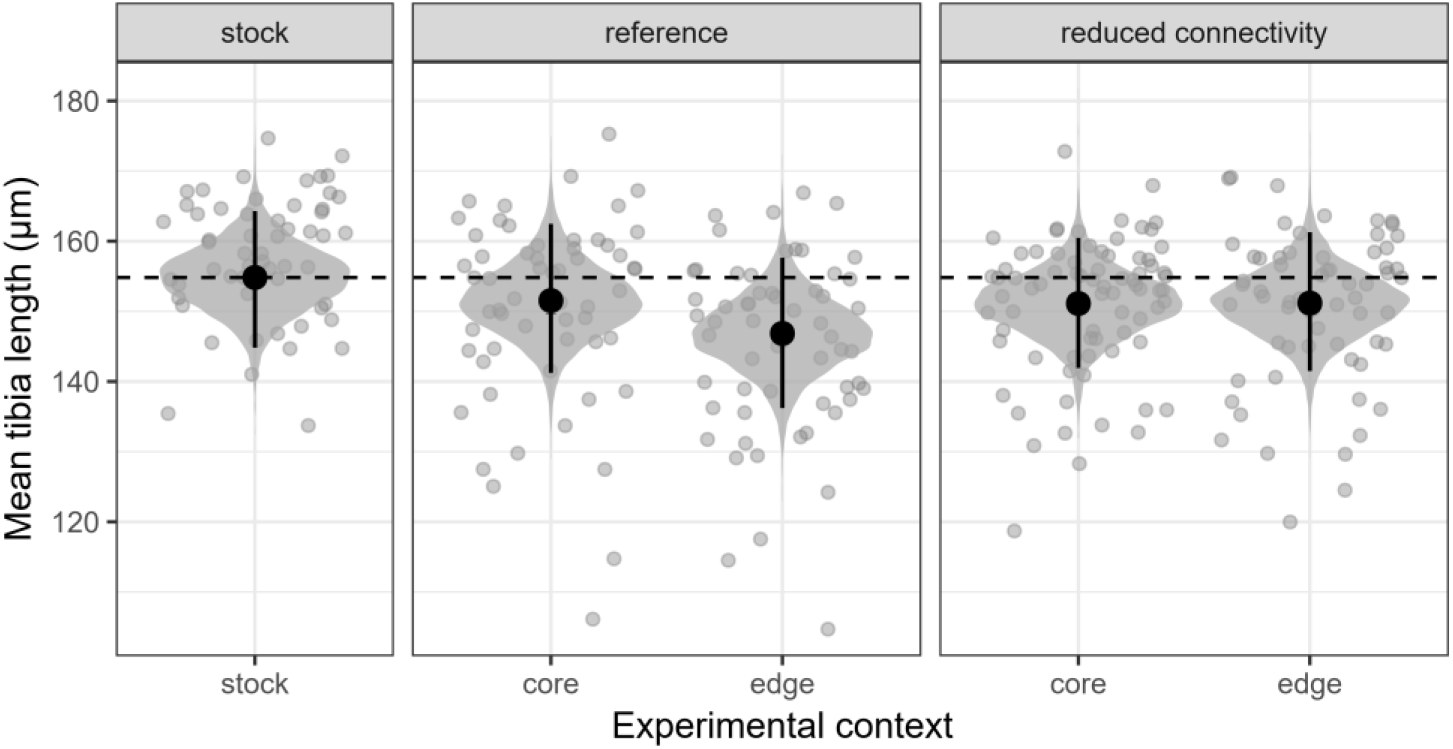
Posterior distribution of mean tibia length (proxy of body size); black dots and segments: posterior means and 95% credible intervals. Grey dots: individual observed values (average of the two observers’ measures). The horizontal dashed line marks the posterior mean for the stocks. See **Supplementary Figure S.2.1** for posterior pairwise comparisons.

We found no evidence that short-term activity had evolved during our experiments (**Fig. 4, Supplementary Figure S.2.2**). Individuals were on average active 53% of the time they were filmed, regardless of connectivity treatments and location (grand mean; 95% CI: [37%; 67%]).

**Figure 4.**
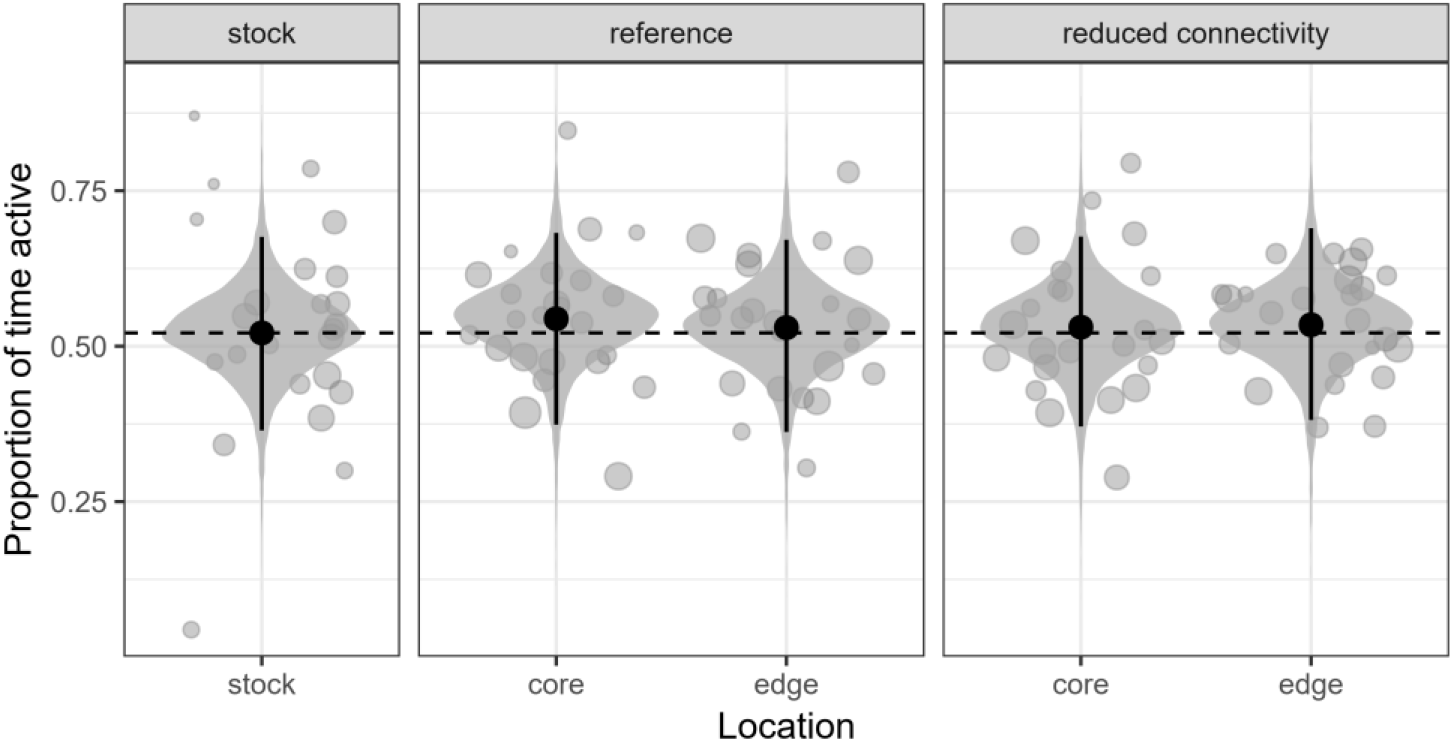
Posterior distributions of mean short-term activity, based on filmed movement tracks; black dots and segments: posterior means and 95% credible intervals. Grey dots: replicate-level observed values; point size is proportional to the total valid observation time for a replicate (sum of all movement bouts). The horizontal dashed line marks the posterior mean for the stocks. See **Supplementary Figure S.2.2** for posterior pairwise comparisons.

We found no consistent deviations from stock population dispersal in the first dispersal experiment, as posteriors were wide (**Fig. 5A**). Dispersal rates were nonetheless higher in edge than core populations, but only in landscapes with reduced connectivity (log(odds ratio) = 0.88 [0.32; 1.45], **Fig. 5A, Supplementary Figure S.2.3**). In the low-density part of the second experiment, there is similarly no consistent evolution of dispersal away from stock population rates (**Fig. 5B**). Similarly to the first experiment however, dispersal from edge populations was higher than in core populations, but this time only in “reference” landscapes (log(odds ratio) = 1.16 [0.14; 2.19], **Fig. 5B, Supplementary Figure S.2.4**).

**Figure 5.**
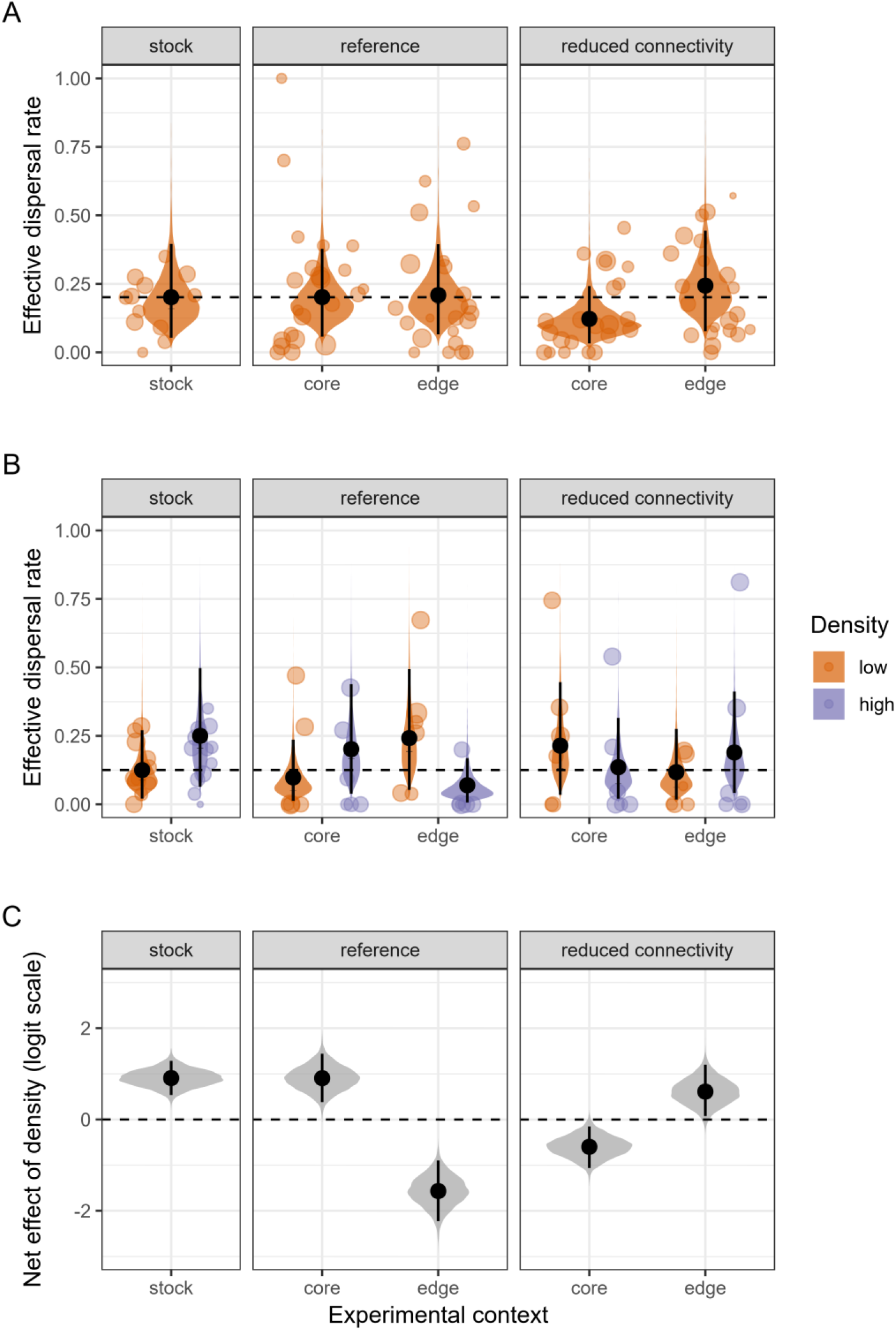
(A; B) Posterior distributions of mean effective dispersal rate, based on either the first experiment (A) or the second experiment (B; one generation later, with some wasps tested at high density). The effect of total fecundity (see **Methods**) on posterior means is averaged out. Black dots and segments: posterior means and 95% credible intervals; the effect of total fecundity (see Methods) on posterior predictions has been averaged out. Coloured dots are observed values, dot size is proportional to total fecundity in each replicate (departure + arrival patches combined). The horizontal dashed lines mark the posterior (low-density) means for the stocks. (C) Net effect of juvenile density on dispersal (difference between posterior mean dispersal at high and low densities, expressed on the logit scale). The horizontal dashed line marks the absence of density-dependence. See **Supplementary Figures S.2.3** and **S.2.4** for the other posterior pairwise comparisons.

Stock populations exhibited positive density-dependent dispersal (log(odds ratio) = 0.91 [0.54; 1.28], **Fig. 5C**). After experimental evolution, this pattern was reversed, leading to negative density-dependent dispersal, in two cases: in wasps coming from edge populations of “reference” landscapes (log(odds ratio) = -1.57 [-2.23; -0.90]) and in wasps from core populations of landscapes with reduced connectivity (log(odds ratio) = -0.60 [-1.06; -0.15])(**Fig. 5C**). Dispersal remained positive density-dependent in the other two connectivity × location treatments (**Fig. 5C**).

Regarding individual fecundity, we found no evidence that landscape connectivity or patch location had any effect in the first fecundity experiment (**Fig. 6A, Supplementary Figure S.2.5**). Similarly, when looking at low-density fecundity in the second (density-dependent) experiment, most of the treatments are not different from each other (**Fig. 6B, Supplementary Figure S.2.6**). The only exception was that low-density edge populations were less fecund than the corresponding stock (log(fold change) = -0.29 [-0.56; -0.02], **Fig. 6B, Supplementary Figure S.2.6**). Additionally, fecundity was not different between low-density and high-density stock populations (**Fig. 6C**); after experimental evolution however, individuals from core populations were less fecund if they came from high-density than if they came from a low-density background, independently of connectivity treatment (log(fold change) = -0.33 [-0.74; 0.02] in reference landscapes, -0.35 [-0.61 -0.11] in landscapes with reduced connectivity, **Fig. 6C**). There was no density effect for individuals from edge populations (**Fig. 6C**). As a consequence of the effects described above, when reared at high densities, wasps coming from the experimental landscapes were in almost all scenarios less fecund on average than the corresponding stock wasps (the exception being wasps from the expansion edge of “reduced connectivity” landscapes; **Fig. 6B, Supplementary Figure S.2.6**).

**Figure 6.**
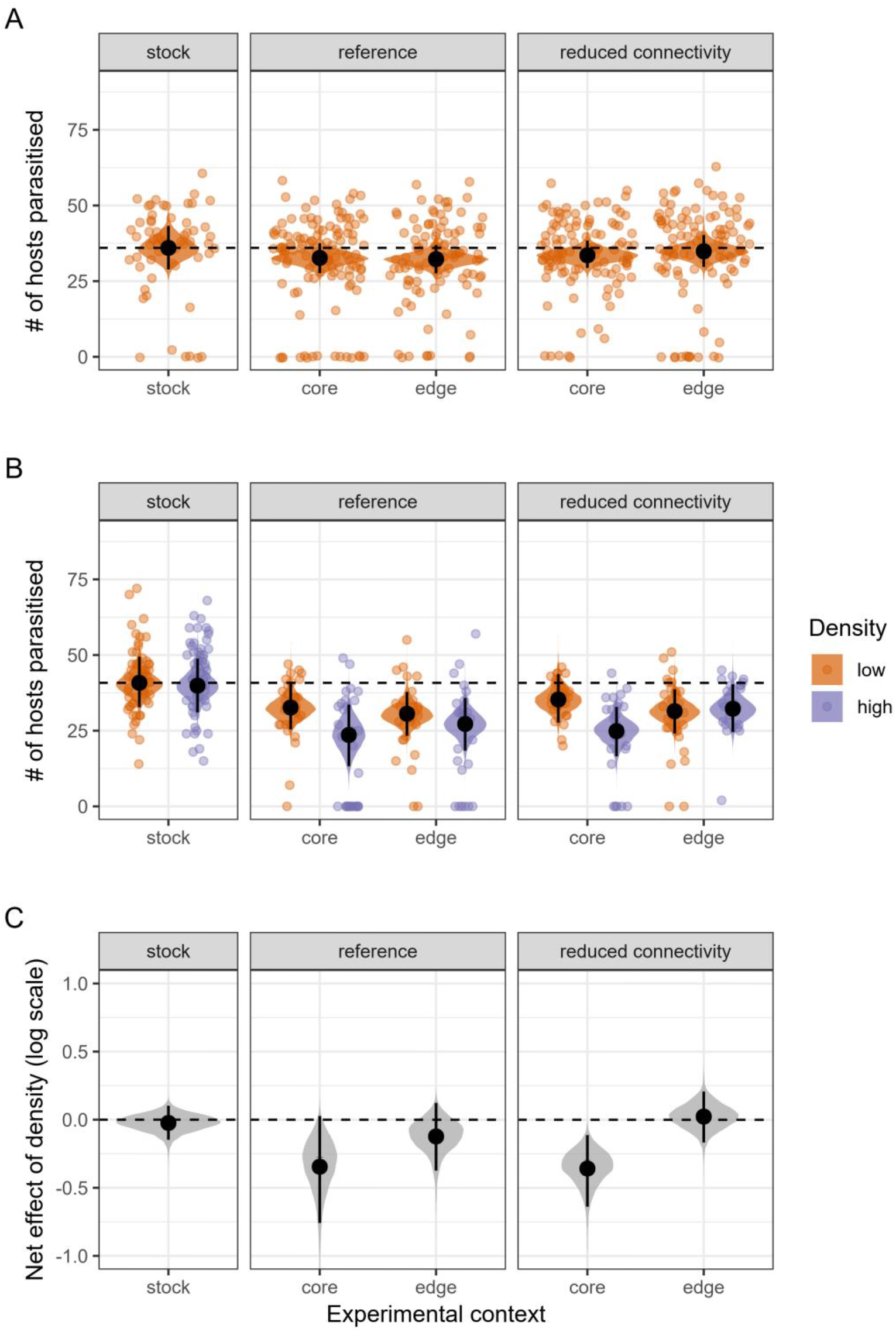
(A; B) Posterior distributions of mean per-capita fecundity, based on either the first experiment (A) or the second experiment (B; one generation later, with some wasps tested at high density). Black dots and segments: posterior means and 95% credible intervals; coloured dots: observed values. The horizontal dashed lines mark the posterior (low-density) means for the stocks. (C) Net effect of juvenile density on fecundity (difference between posterior mean fecundity at high and low densities, expressed on the log scale). The horizontal dashed line marks the absence of density-dependence. See **Supplementary Figures S.2.5** and **S.2.6** for the other posterior pairwise comparisons.

## Discussion

We showed previously that variation in landscape connectivity shapes the position of experimental range expansions on the pushed/pulled expansion axis in *Trichogramma* wasps (Dahirel et al., 2021). We here find that these previously documented changes in expansion and neutral diversity dynamics due to connectivity were not consistently accompanied by clear phenotypic shifts at the range edge. However, we found some indications that the density-dependence of dispersal, one of the two key parameters determining the pushed vs. pulled nature of expansions (Birzu et al., 2019), may change during the range expansion process, and these changes seemed to depend on the connectivity level.

We did not find any clear evidence for evolutionary changes in size or short-term activity, nor in fecundity or effective dispersal when density-dependence was ignored (**Figs 3 to 6**). Reproductive success did vary between treatments, but only in the density-dependent experiment, and the only consistent shift was that at high densities, post-experimental evolution wasps were less fecund than wasps from stock populations (irrespective of connectivity or patch location)(**Fig. 6**). We found some evidence of higher low-density dispersal in edge compared to core populations, as expected from theory (Chuang & Peterson, 2016; Shine et al., 2011; Travis et al., 2009). However, our experiments are here inconsistent: low-density dispersal was higher in edge vs. core patches only in “reduced connectivity” landscapes in one dispersal experiment, and only in “reference” landscapes in the other dispersal experiment (**Fig. 5, Supplementary Figures S.2.3** and **S.2.4**). There was also no clear divergence from the starting stock populations themselves (**Fig. 5, Supplementary Figures S.2.3** and **S.2.4**). Our results here contrast previous experiments (e.g. Williams, Kendall, et al., 2016) and theoretical models (Williams, Snyder, et al., 2016) that showed both evolutionary changes in key traits along expansion edges, and that this evolution was accelerated in more fragmented environments. While clear increases in average dispersal or per capita growth rates are often expected at the edge of range expansions (Chuang & Peterson, 2016; Fronhofer et al., 2017; Phillips & Perkins, 2019; Van Petegem et al., 2018; Weiss-Lehman et al., 2017), there are enough exceptions to the “rule” (Chuang & Peterson, 2016; Van Petegem et al., 2018; Wolz et al., 2020) for these null/uncertain results not to be entirely surprising by themselves. Trade-offs among traits may provide a mechanistic explanation for this absence of evolutionary response: Williams et al. (2016) and Urquhart and Williams (2021) showed that the shape and strength of trade-offs among traits may influence whether the way these traits evolve during expansion is sensitive to connectivity. Similarly, Ochocki et al. (2020) showed, using simulations, that genetic trade-offs between dispersal and fecundity may reduce, and in some cases prevent, the evolution of these traits at the range edge. As Ochocki et al. (2020) mentioned, knowledge about trait architecture may be key to interpret eco-evolutionary outcomes of range expansion, and the variability among species/studies.

Irrespective of the impact of trade-offs, focusing on trait(s) expression at only one density is limiting, as the density-dependence of dispersal or growth actually plays a key role in shaping the dynamics of range expansions (Birzu et al., 2019). We previously found that, in our *Trichogramma* experimental system, expansions were more pushed when connectivity was reduced, which means that connectivity influenced the density-dependence of growth and/or dispersal, through plastic and/or evolved responses (Dahirel et al., 2021). While our data are limited (see below), we here find some evidence for density-dependent effective dispersal and reproductive success, and for variation in this density-dependence across landscape connectivity contexts. Because we tested wasps using a common garden protocol, the differences we observed are likely the result of evolutionary divergence during expansions (although parental and grandparental effects on density-dependent dispersal cannot be ruled out entirely; Bitume et al., 2014).

First, in core populations, the experiments showed a link between density and per capita fecundity that is absent from edge populations (as well as from stock populations). Specifically, wasps coming from these core lineages had fewer offspring on average when raised in high-density conditions (**Fig. 6**). This lower fecundity is expected if there is an egg number-egg size trade-off, as higher competition in core patches would favour larger, more competitive larvae (Segoli & Wajnberg, 2020). For instance, in *Callosobruchus chinensis* beetles parasitising seeds, higher larval competition within seeds leads to adults producing both a reduced number of eggs (Vamosi, 2005) and larger eggs (after accounting for emerging female size; Yanagi et al., 2013). Alternatively, core populations may have evolved a higher propensity to superparasitism, since there individuals experienced higher densities, and encounters with hosts parasitized by other wasps, more frequently (Van Alphen & Visser, 1990). Wasps emerging from superparasitized hosts tend to be smaller and less fecund (Durocher-Granger et al., 2011). To confirm and disentangle these hypotheses however, further experiments would be needed to determine whether there is actually an egg number-egg size trade-off in our tested populations.

Second, *Trichogramma* wasps from the stock populations dispersed more on average if they came from a high density background (**Fig. 5**). This finding fits with the classic view of density-dependent dispersal as a response to increased competition (Bowler & Benton, 2005; Harman et al., 2020), and is a logical extension of previous results showing *Trichogramma brassicae* wasps left host eggs patches earlier if more were already parasitized (Wajnberg et al., 2000). The direction of this density-dispersal relationship was reversed in “reference” edge populations after 12 generations of evolution and expansion (**Fig. 5**), with wasps dispersing more from low-density populations. Our results here broadly agree with theory, which tends to predict the loss of positive density-dependent dispersal at low-density expansion edges (cf e.g. Travis et al., 2009). There is one key nuance in that theoretical models often predict unconditional high dispersal over most of the range of densities as a result, where we found a shift to negative density-dependent dispersal. However, it is difficult to say whether the former is the “true” expected endpoint during range expansions, given many dispersal models are designed or parameterized in a way that excludes the possibility of negative density-dependent dispersal (Kun & Scheuring, 2006; e.g. Poethke & Hovestadt, 2002; Travis et al., 2009). Indeed, other empirical studies show that shifts to negative density-dependent dispersal can happen at the edge of range expansions (Fronhofer et al., 2017; Simmons & Thomas, 2004). In any case, the key result remains consistent with theory, in that evolution at the range edge removes the positive density-dependence of dispersal that existed initially. By contrast, when connectivity was reduced, no clear evolutionary changes in dispersal reaction norm occurred at the range edge (**Fig. 5**): the slope remained positive, albeit slightly shallower (as in Weiss-Lehman et al., 2017). Our results at the expanding edge are here consistent with existing theory, since strong enough increases in dispersal costs (such as those that may be caused by reduced connectivity) are predicted to favour more positive density-dependent dispersal (Govindan et al., 2015; Rodrigues & Johnstone, 2014; Travis et al., 1999). However, in core populations dispersal actually became negative density-dependent when connectivity was reduced (**Fig. 5**), seemingly contradicting the previous argument. As discussed above, the theory explaining negative density-dependent dispersal is much less developed in stable metapopulations, let alone in range expansions. Among the few existing models, Rodrigues and Johnstone (2014) predicted that, at least in a non-expanding context, reduced temporal variability should favour negative density-dependent dispersal. Reusing population size data in Dahirel et al. (2021), we find that reduced connectivity did indeed lead to lower temporal variability in core patches (**Supplementary Material S3**). We can tentatively interpret our results as the interplay of three “forces.” On one side, the expansion process itself drives the loss of positive density-dependent dispersal at the expansion edge. On the other side, connectivity has dual and contradictory effects: the direct effects of reduced connectivity on dispersal costs would favour positive density-dependent dispersal; while the indirect effects through demographic stochasticity would favour negative density-dependent dispersal.

Taken altogether, our results confirm the importance of context-dependence when studying dispersal (Bonte & Dahirel, 2017; Matthysen, 2012). This is especially true for range expansions, which are often associated with a core-to-edge density gradient. We argue that not considering this context-dependence may explain (some of the) previous failures to detect trait evolution during range expansions (see e.g. compilation in Chuang & Peterson, 2016), and we recommend testing for density-dependence whenever it is logistically possible (as in e.g. Weiss-Lehman et al., 2017).

We acknowledge that these findings regarding dispersal come from the experiment with the lowest sample size within this study (see **Methods** and **Fig. 2**) and need further confirmation. High numbers of replicate landscapes in experimental (and natural) expansion studies are especially important if we want to make generalizable inferences and predictions, due to the key role of evolutionary stochasticity in shaping outcomes (Phillips, 2015; Weiss-Lehman et al., 2017; Williams et al., 2019). Moreover, we only sampled a limited subset of this species genetic diversity, and the three strains we work with may be biased towards some life histories. Further comparative analyses would be important to determine the effect of initial genetic/phenotypic variation on ecological and evolutionary dynamics during expansions (Miller et al., 2020). Finally, the fact we only detected evolutionary changes in the density-dependent experiment may be because we used, due again to limited sample size for some traits, a coarse definition of “core” vs. “edge” patches that ignored variation in distances travelled since the start of expansions/expansion speed. Despite these potential issues, our findings on the evolution of density-dependent dispersal are fully consistent with previous results and expectations regarding pushed vs. pulled expansions (Birzu et al., 2019; Dahirel et al., 2021), as detailed below. As such, we see them as a first step towards research that better accounts for the complexities of eco-evolutionary dynamics during (pushed) range expansions, and hope that they encourage further studies on the subject.

## Conclusion: implications for the evolution of pushed expansion

Although *Trichogramma brassicae* wasps start the experiments with positive density-dependent dispersal, it seems edge populations evolve away from that strategy rapidly if left to expand in relatively well connected “reference” landscapes (**Fig. 6**). Our experimental results agree with Erm and Phillips (2020)’s model, in which evolution should lead initially pushed expansions to become pulled (in their case with Allee effect-induced pushed expansions, in ours with density-dependent dispersal). The fundamental mechanism is the same in both cases: low densities at the expanding range edge mean that anything that disperses or grows worse at low densities will be outperformed/outrun, leading to an accumulation of individuals that disperse/grow well at low densities. Taken alone, these results would imply pushed range expansions are rare in nature since evolution would tend to “erase” them, or at least not as common as would be expected from general frequencies of Allee effects (Gregory et al., 2010) and positive density-dependent dispersal (Harman et al., 2020) in non-expanding populations. On the other hand, we found that positive density-dependent dispersal is comparatively maintained in edge populations, even after >10 generations of expansion, in landscapes with reduced connectivity. Accordingly, these expansions were previously shown to have more “pushed” characteristics than controls (Dahirel et al., 2021). Thus, persistent pushed expansions may actually be favoured in the many landscapes experiencing anthropogenic connectivity loss. In any case, our results show that environmental conditions and constraints may be key to the maintenance of pushed expansion dynamics in the face of evolutionary dynamics, and that the context dependence of pushed expansions needs to be further explored. We note however that more work (experimental or modelling) is needed to confirm this, especially to understand the implications of our results on longer time scales (Birzu et al., 2019).

Pushed and pulled expansions can differ in (relative) speed, genetic diversity (Dahirel et al., 2021) and, as our results show here, phenotypic composition. Lineages/individuals with different dispersal strategies may also differ in traits influencing population stability (Jacob et al., 2019) or ecosystem functioning (Cote et al., 2017; Little et al., 2019). Understanding what environmental conditions favour or disfavour the evolutionary maintenance of “pushiness” during expansions may help more generally to understand the evolution of many traits during range expansions, and the possible functional effects of expanding species on ecosystems.

## Data accessibility

Data and R scripts to reproduce all analyses presented in this paper are available on Github (https://github.com/mdahirel/pushed-pulled-2020-phenotype) and archived in Zenodo (v1.2; https://doi.org/10.5281/zenodo.4570235).

## Supplementary material

Additional supplementary material and analyses are available online from the same sources as above.

## Acknowledgements

We thank participants to the 2019 conference of the British Ecological Society, as well as Thomas Guillemaud, for their questions during talks and discussions leading up to this paper. We also thank Inês Fragata and three anonymous reviewers for helpful comments on a previous version of this paper. Version 4 of this preprint has been peer-reviewed and recommended by Peer Community In Evolutionary Biology (https://doi.org/10.24072/pci.evolbiol.100133).

## References

Allan, C., Burel, J.-M., Moore, J., Blackburn, C., Linkert, M., Loynton, S., MacDonald, D., Moore, W. J., Neves, C., Patterson, A., Porter, M., Tarkowska, A., Loranger, B., Avondo, J., Lagerstedt, I., Lianas, L., Leo, S., Hands, K., Hay, R. T., … Swedlow, J. R. (2012). OMERO: flexible, model-driven data management for experimental biology. Nature Methods, 9(3), 245–253. https://doi.org/10.1038/nmeth.1896

Allee, W. C., & Bowen, E. S. (1932). Studies in animal aggregations: Mass protection against colloidal silver among goldfishes. Journal of Experimental Zoology, 61(2), 185–207. https://doi.org/10.1002/jez.1400610202

Benhamou, S. (2004). How to reliably estimate the tortuosity of an animal’s path. Journal of Theoretical Biology, 229(2), 209–220. https://doi.org/10.1016/j.jtbi.2004.03.016

Birzu, G., Hallatschek, O., & Korolev, K. S. (2018). Fluctuations uncover a distinct class of traveling waves. Proceedings of the National Academy of Sciences, 115(16), E3645–E3654. https://doi.org/10.1073/pnas.1715737115

Birzu, G., Matin, S., Hallatschek, O., & Korolev, K. S. (2019). Genetic drift in range expansions is very sensitive to density dependence in dispersal and growth. Ecology Letters, 22(11), 1817–1827. https://doi.org/10.1111/ele.13364

Bitume, E. V., Bonte, D., Ronce, O., Olivieri, I., & Nieberding, C. M. (2014). Dispersal distance is influenced by parental and grand-parental density. Proceedings of the Royal Society of London B: Biological Sciences, 281(1790), 20141061. https://doi.org/10.1098/rspb.2014.1061

Bonte, D., & Dahirel, M. (2017). Dispersal: a central and independent trait in life history. Oikos, 126(4), 472–479. https://doi.org/10.1111/oik.03801

Bowler, D. E., & Benton, T. G. (2005). Causes and consequences of animal dispersal strategies: relating individual behaviour to spatial dynamics. Biological Reviews, 80(2), 205–225. https://doi.org/10.1017/S1464793104006645

Branson, K., Robie, A. A., Bender, J., Perona, P., & Dickinson, M. H. (2009). High-throughput ethomics in large groups of Drosophila. Nature Methods, 6(6), 451–457. https://doi.org/10.1038/nmeth.1328

Burton, O. J., Phillips, B. L., & Travis, J. M. J. (2010). Trade-offs and the evolution of life-histories during range expansion. Ecology Letters, 13(10), 1210–1220. https://doi.org/10.1111/j.1461-0248.2010.01505.x

Bürkner, P.-C. (2017). brms: an R package for Bayesian multilevel models using Stan. Journal of Statistical Software, 80(1), 1–28. https://doi.org/10.18637/jss.v080.i01

Carpenter, B., Gelman, A., Hoffman, M. D., Lee, D., Goodrich, B., Betancourt, M., Brubaker, M., Guo, J., Li, P., & Riddell, A. (2017). Stan: a probabilistic programming language. Journal of Statistical Software, 76(1), 1–32. https://doi.org/10.18637/jss.v076.i01

Chuang, A., & Peterson, C. R. (2016). Expanding population edges: theories, traits, and trade-offs. Global Change Biology, 22(2), 494–512. https://doi.org/10.1111/gcb.13107

Clutton-Brock, T. H., & Harvey, P. H. (1977). Primate ecology and social organization. Journal of Zoology, 183(1), 1–39. https://doi.org/10.1111/j.1469-7998.1977.tb04171.x

Corrigan, J. E., Laing, J. E., & Zubricky, J. S. (1995). Effects of parasitoid to host ratio and time of day of parasitism on development and emergence of Trichogramma minutum (Hymenoptera: Trichogrammatidae) parasitizing eggs of Ephestia kuehniella (Lepidoptera: Pyralidae). Annals of the Entomological Society of America, 88(6), 773–780. https://doi.org/10.1093/aesa/88.6.773

Cote, J., Brodin, T., Fogarty, S., & Sih, A. (2017). Non-random dispersal mediates invader impacts on the invertebrate community. Journal of Animal Ecology, 86(6), 1298–1307. https://doi.org/10.1111/1365-2656.12734

Courchamp, F., Berec, L., & Gascoigne, J. (2008). Allee effects in ecology and conservation. Oxford University Press.

Cwynar, L. C., & MacDonald, G. M. (1987). Geographical variation of lodgepole pine in relation to population history. The American Naturalist, 129(3), 463–469. https://doi.org/10.1086/284651

Dahirel, M., Bertin, A., Haond, M., Blin, A., Lombaert, E., Calcagno, V., Fellous, S., Mailleret, L., Malausa, T., & Vercken, E. (2021). Shifts from pulled to pushed range expansions caused by reduction of landscape connectivity. Oikos, 130(5), 708–724. https://doi.org/10.1111/oik.08278

Deforet, M., Carmona-Fontaine, C., Korolev, K. S., & Xavier, J. B. (2019). Evolution at the edge of expanding populations. The American Naturalist, 194(3), 291–305. https://doi.org/10.1086/704594

Des Roches, S., Post, D. M., Turley, N. E., Bailey, J. K., Hendry, A. P., Kinnison, M. T., Schweitzer, J. A., & Palkovacs, E. P. (2018). The ecological importance of intraspecific variation. Nature Ecology & Evolution, 2(1), 57–64. https://doi.org/10.1038/s41559-017-0402-5

Durocher-Granger, L., Martel, V., & Boivin, G. (2011). Gamete number and size correlate with adult size in the egg parasitoid Trichogramma euproctidis. Entomologia Experimentalis Et Applicata, 140(3), 262–268. https://doi.org/10.1111/j.1570-7458.2011.01158.x

Erm, P., & Phillips, B. L. (2020). Evolution transforms pushed waves into pulled waves. The American Naturalist, 195(3), E87–E99. https://doi.org/10.1086/707324

Fellous, S., Angot, G., Orsucci, M., Migeon, A., Auger, P., Olivieri, I., & Navajas, M. (2014). Combining experimental evolution and field population assays to study the evolution of host range breadth. Journal of Evolutionary Biology, 27(5), 911–919. https://doi.org/10.1111/jeb.12362

Fronhofer, E. A., Gut, S., & Altermatt, F. (2017). Evolution of density-dependent movement during experimental range expansions. Journal of Evolutionary Biology, 30(12), 2165–2176. https://doi.org/10.1111/jeb.13182

Fronhofer, E. A., Legrand, D., Altermatt, F., Ansart, A., Blanchet, S., Bonte, D., Chaine, A., Dahirel, M., Laender, F. D., Raedt, J. D., di Gesu, L., Jacob, S., Kaltz, O., Laurent, E., Little, C. J., Madec, L., Manzi, F., Masier, S., Pellerin, F., … Cote, J. (2018). Bottom-up and top-down control of dispersal across major organismal groups. Nature Ecology & Evolution, 2(12), 1859–1863. https://doi.org/10.1038/s41559-018-0686-0

Gabry, J., Simpson, D., Vehtari, A., Betancourt, M., & Gelman, A. (2019). Visualization in Bayesian workflow. Journal of the Royal Statistical Society: Series A (Statistics in Society), 182(2), 389–402. https://doi.org/10.1111/rssa.12378

Gandhi, S. R., Korolev, K. S., & Gore, J. (2019). Cooperation mitigates diversity loss in a spatially expanding microbial population. Proceedings of the National Academy of Sciences, 116(47), 23582–23587. https://doi.org/10.1073/pnas.1910075116

Govindan, B. N., Feng, Z., DeWoody, Y. D., & Swihart, R. K. (2015). Intermediate disturbance in experimental landscapes improves persistence of beetle metapopulations. Ecology, 96(3), 728–736. https://doi.org/10.1890/14-0044.1

Gralka, M., & Hallatschek, O. (2019). Environmental heterogeneity can tip the population genetics of range expansions. eLife, 8, e44359. https://doi.org/10.7554/eLife.44359

Gregory, S. D., Bradshaw, C. J. A., Brook, B. W., & Courchamp, F. (2010). Limited evidence for the demographic Allee effect from numerous species across taxa. Ecology, 91(7), 2151–2161. https://doi.org/10.1890/09-1128.1

Hadfield, J. D., & Nakagawa, S. (2010). General quantitative genetic methods for comparative biology: phylogenies, taxonomies and multi-trait models for continuous and categorical characters. Journal of Evolutionary Biology, 23(3), 494–508. https://doi.org/10.1111/j.1420-9101.2009.01915.x

Harman, R. R., Goddard, J., Shivaji, R., & Cronin, J. T. (2020). Frequency of occurrence and population-dynamic consequences of different forms of density-dependent emigration. The American Naturalist, 195(5), 851–867. https://doi.org/10.1086/708156

Hunter, M., Krishnan, N., Liu, T., Möbius, W., & Fusco, D. (2021). Virus-host interactions shape viral dispersal giving rise to distinct classes of traveling waves in spatial expansions. Physical Review X, 11(2), 021066. https://doi.org/10.1103/PhysRevX.11.021066

Jacob, S., Chaine, A. S., Huet, M., Clobert, J., & Legrand, D. (2019). Variability in dispersal syndromes is a key driver of metapopulation dynamics in experimental microcosms. The American Naturalist, 194(5), 613–626. https://doi.org/10.1086/705410

Kay, M. (2019). tidybayes: tidy data and geoms for Bayesian models. https://doi.org/10.5281/zenodo.1308151

Kun, Á., & Scheuring, I. (2006). The evolution of density-dependent dispersal in a noisy spatial population model. Oikos, 115(2), 308–320. https://doi.org/10.1111/j.2006.0030-1299.15061.x

Lenoir, J., Bertrand, R., Comte, L., Bourgeaud, L., Hattab, T., Murienne, J., & Grenouillet, G. (2020). Species better track climate warming in the oceans than on land. Nature Ecology & Evolution. https://doi.org/10.1038/s41559-020-1198-2

Lewis, M., Petrovskii, S. V., & Potts, J. (2016). The mathematics behind biological invasions. Springer International Publishing. https://doi.org/10.1007/978-3-319-32043-4

Little, C. J., Fronhofer, E. A., & Altermatt, F. (2019). Dispersal syndromes can impact ecosystem functioning in spatially structured freshwater populations. Biology Letters, 15(3), 20180865. https://doi.org/10.1098/rsbl.2018.0865

Lutscher, F., & Musgrave, J. A. (2017). Behavioral responses to resource heterogeneity can accelerate biological invasions. Ecology, 98(5), 1229–1238. https://doi.org/10.1002/ecy.1773

Matthysen, E. (2012). Multicausality of dispersal: a review. In J. Clobert, M. Baguette, T. G. Benton, & J. M. Bullock (Eds.), Dispersal ecology and evolution (pp. 3–18). Oxford University Press.

Matthysen, E. (2005). Density-dependent dispersal in birds and mammals. Ecography, 28(3), 403–416. https://doi.org/10.1111/j.0906-7590.2005.04073.x

McElreath, R. (2020). Statistical rethinking: a Bayesian course with examples in R and Stan (2nd edition). Chapman and Hall/CRC.

McLean, D. J., & Volponi, M. A. S. (2018). trajr: An R package for characterisation of animal trajectories. Ethology, 124(6), 440–448. https://doi.org/10.1111/eth.12739

Merwin, A. C. (2019). Flight capacity increases then declines from the core to the margins of an invasive species’ range. Biology Letters, 15(11), 20190496. https://doi.org/10.1098/rsbl.2019.0496

Miller, T. E. X., Angert, A. L., Brown, C. D., Lee-Yaw, J. A., Lewis, M., Lutscher, F., Marculis, N. G., Melbourne, B. A., Shaw, A. K., Szűcs, M., Tabares, O., Usui, T., Weiss-Lehman, C., & Williams, J. L. (2020). Eco-evolutionary dynamics of range expansion. Ecology, n/a(n/a), e03139. https://doi.org/10.1002/ecy.3139

Mishra, A., Chakraborty, P. P., & Dey, S. (2020). Dispersal evolution diminishes the negative density dependence in dispersal. Evolution, evo.14070. https://doi.org/10.1111/evo.14070

Morel-Journel, T., Girod, P., Mailleret, L., Auguste, A., Blin, A., & Vercken, E. (2016). The highs and lows of dispersal: how connectivity and initial population size jointly shape establishment dynamics in discrete landscapes. Oikos, 125(6), 769–777. https://doi.org/10.1111/oik.02718

Ochocki, B. M., Saltz, J. B., & Miller, T. E. X. (2020). Demography-dispersal trait correlations modify the ecoevolutionary dynamics of range expansion. The American Naturalist, 195(2), 231–246. https://doi.org/10.1086/706904

Pachepsky, E., & Levine Jonathan M. (2011). Density dependence slows invader spread in fragmented landscapes. The American Naturalist, 177(1), 18–28. https://doi.org/10.1086/657438

Pedersen, T. L. (2019). patchwork: the composer of plots. https://CRAN.R-project.org/package=patchwork

Perkins, T. A., Phillips, B. L., Baskett, M. L., & Hastings, A. (2013). Evolution of dispersal and life history interact to drive accelerating spread of an invasive species. Ecology Letters, 16(8), 1079–1087. https://doi.org/10.1111/ele.12136

Phillips, B. L. (2015). Evolutionary processes make invasion speed difficult to predict. Biological Invasions, 17(7), 1949–1960. https://doi.org/10.1007/s10530-015-0849-8

Phillips, B. L., & Perkins, T. A. (2019). Spatial sorting as the spatial analogue of natural selection. Theoretical Ecology. https://doi.org/10.1007/s12080-019-0412-9

Poethke, H. J., & Hovestadt, T. (2002). Evolution of density–and patch–size–dependent dispersal rates. Proceedings of the Royal Society of London B: Biological Sciences, 269(1491), 637–645. https://doi.org/10.1098/rspb.2001.1936

R Core Team. (2021). R: a language and environment for statistical computing (Version 4.0.4) [Computer software]. R Foundation for Statistical Computing. https://www.R-project.org/

Raffard, A., Santoul, F., Cucherousset, J., & Blanchet, S. (2019). The community and ecosystem consequences of intraspecific diversity: a meta-analysis. Biological Reviews, 94(2), 648–661. https://doi.org/10.1111/brv.12472

Renault, D., Laparie, M., McCauley, S. J., & Bonte, D. (2018). Environmental adaptations, ecological filtering, and dispersal central to insect invasions. Annual Review of Entomology, 63(1), 345–368. https://doi.org/10.1146/annurev-ento-020117-043315

Rodrigues, A. M. M., & Johnstone, R. A. (2014). Evolution of positive and negative density-dependent dispersal. Proceedings of the Royal Society of London B: Biological Sciences, 281(1791), 20141226. https://doi.org/10.1098/rspb.2014.1226

Ronce, O., & Clobert, J. (2012). Dispersal syndromes. In J. Clobert, M. Baguette, T. G. Benton, & J. M. Bullock (Eds.), Dispersal ecology and evolution (pp. 119–138). Oxford University Press.

Roques, L., Garnier, J., Hamel, F., & Klein, E. K. (2012). Allee effect promotes diversity in traveling waves of colonization. Proceedings of the National Academy of Sciences, 109(23), 8828–8833. https://doi.org/10.1073/pnas.1201695109

Schreiber, S. J., & Beckman, N. G. (2020). Individual variation in dispersal and fecundity increases rates of spatial spread. AoB PLANTS, 12(3), Article 3. https://doi.org/10.1093/aobpla/plaa001

Segoli, M., & Wajnberg, E. (2020). The combined effect of host and food availability on optimized parasitoid life-history traits based on a three-dimensional trade-off surface. Journal of Evolutionary Biology, 33(6), 850–857. https://doi.org/10.1111/jeb.13617

Shine, R., Brown, G. P., & Phillips, B. L. (2011). An evolutionary process that assembles phenotypes through space rather than through time. Proceedings of the National Academy of Sciences, 108(14), 5708–5711. https://doi.org/10.1073/pnas.1018989108

Sibly, R. M., & Hone, J. (2002). Population growth rate and its determinants: an overview. Philosophical Transactions of the Royal Society of London. Series B: Biological Sciences, 357(1425), 1153–1170. https://doi.org/10.1098/rstb.2002.1117

Simmons, Adam D., & Thomas, Chris D. (2004). Changes in dispersal during species’ range expansions. The American Naturalist, 164(3), 378–395. https://doi.org/10.1086/423430

Stokes, A. N. (1976). On two types of moving front in quasilinear diffusion. Mathematical Biosciences, 31(3), 307–315. https://doi.org/10.1016/0025-5564(76)90087-0

St-Onge, M., Cormier, D., Todorova, S., & Lucas, É. (2014). Comparison of Ephestia kuehniella eggs sterilization methods for Trichogramma rearing. Biological Control, 70, 73–77. https://doi.org/10.1016/j.biocontrol.2013.12.006

Travis, J. M., & Dytham, C. (2002). Dispersal evolution during invasions. Evolutionary Ecology Research, 4(8), 1119–1129. http://www.evolutionary-ecology.com/abstracts/v04/1413.html

Travis, J. M. J., Murrell, D. J., & Dytham, C. (1999). The evolution of density–dependent dispersal. Proceedings of the Royal Society of London B: Biological Sciences, 266(1431), 1837–1842. https://doi.org/10.1098/rspb.1999.0854

Travis, J. M. J., Mustin, K., Benton, T. G., & Dytham, C. (2009). Accelerating invasion rates result from the evolution of density-dependent dispersal. Journal of Theoretical Biology, 259(1), 151–158. https://doi.org/10.1016/j.jtbi.2009.03.008

Urquhart, C. A., & Williams, J. L. (2021). Trait correlations and landscape fragmentation jointly alter expansion speed via evolution at the leading edge in simulated range expansions. Theoretical Ecology. https://doi.org/10.1007/s12080-021-00503-z

Vamosi, S. M. (2005). Interactive effects of larval host and competition on adult fitness: an experimental test with seed beetles (Coleoptera: Bruchidae). Functional Ecology, 19(5), 859–864. https://doi.org/10.1111/j.1365-2435.2005.01029.x

Van Alphen, J. J. M., & Visser, M. E. (1990). Superparasitism as an adaptive strategy for insect parasitoids. Annual Review of Entomology, 35(1), 59–79. https://doi.org/10.1146/annurev.en.35.010190.000423

Van Petegem, K., Moerman, F., Dahirel, M., Fronhofer, E. A., Vandegehuchte, M. L., Van Leeuwen, T., Wybouw, N., Stoks, R., & Bonte, D. (2018). Kin competition accelerates experimental range expansion in an arthropod herbivore. Ecology Letters, 21(2), 225–234. https://doi.org/10.1111/ele.12887

Vehtari, A., Gelman, A., Simpson, D., Carpenter, B., & Bürkner, P.-C. (2020). Rank-normalization, folding, and localization: an improved 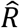 for assessing convergence of MCMC. Bayesian Analysis. https://doi.org/10.1214/20-BA1221

Violle, C., Enquist, B. J., McGill, B. J., Jiang, L., Albert, C. H., Hulshof, C., Jung, V., & Messier, J. (2012). The return of the variance: intraspecific variability in community ecology. Trends in Ecology & Evolution, 27(4), 244–252. https://doi.org/10.1016/j.tree.2011.11.014

Wajnberg, E., Fauvergue, X., & Pons, O. (2000). Patch leaving decision rules and the Marginal Value Theorem: an experimental analysis and a simulation model. Behavioral Ecology, 11(6), 577–586. https://doi.org/10.1093/beheco/11.6.577

Weiss-Lehman, C., Hufbauer, R. A., & Melbourne, B. A. (2017). Rapid trait evolution drives increased speed and variance in experimental range expansions. Nature Communications, 8, 14303. https://doi.org/10.1038/ncomms14303

Wickham, H., Averick, M., Bryan, J., Chang, W., McGowan, L., François, R., Grolemund, G., Hayes, A., Henry, L., Hester, J., Kuhn, M., Pedersen, T., Miller, E., Bache, S., Müller, K., Ooms, J., Robinson, D., Seidel, D., Spinu, V., … Yutani, H. (2019). Welcome to the Tidyverse. Journal of Open Source Software, 4(43), 1686. https://doi.org/10.21105/joss.01686

Williams, J. L., Hufbauer, R. A., & Miller, T. E. X. (2019). How evolution modifies the variability of range expansion. Trends in Ecology & Evolution, 34(10), 903–913. https://doi.org/10.1016/j.tree.2019.05.012

Williams, J. L., Kendall, B. E., & Levine, J. M. (2016). Rapid evolution accelerates plant population spread in fragmented experimental landscapes. Science, 353(6298), 482–485. https://doi.org/10.1126/science.aaf6268

Williams, J. L., & Levine Jonathan M. (2018). Experimental evidence that density dependence strongly influences plant invasions through fragmented landscapes. Ecology, 99(4), 876–884. https://doi.org/10.1002/ecy.2156

Williams, J. L., Snyder, R. E., & Levine, J. M. (2016). The influence of evolution on population spread through patchy landscapes. The American Naturalist, 188(1), 15–26. https://doi.org/10.1086/686685

Wolz, M., Klockmann, M., Schmitz, T., Pekár, S., Bonte, D., & Uhl, G. (2020). Dispersal and life-history traits in a spider with rapid range expansion. Movement Ecology, 8(1), 2. https://doi.org/10.1186/s40462-019-0182-4

Yanagi, S., Saeki, Y., & Tuda, M. (2013). Adaptive egg size plasticity for larval competition and its limits in the seed beetle Callosobruchus chinensis. Entomologia Experimentalis Et Applicata, 148(2), 182–187. https://doi.org/10.1111/eea.12088

