## Supplementary material for "Landscape connectivity alters the evolution of density-dependent dispersal during pushed range expansions": supplementary.html


Code 

- Download Rmd

### Supplementary Material for: “Landscape connectivity alters the evolution of density-dependent dispersal during pushed range expansions”

###### Maxime Dahirel, Aline Bertin, Vincent Calcagno, Camille Duraj, Simon Fellous, Géraldine Groussier, Eric Lombaert, Ludovic Mailleret, Anaël Marchand, Elodie Vercken

### S.1 - Model descriptions

We outline here the structure of the models presented in the main text, as well as the corresponding priors. Notation conventions and (weakly informative) prior choices mostly follow McElreath (2020). We use the \(\mathrm{Half-Normal}(0,\sigma)\) notation to denote a half-normal distribution based on a \(\mathrm{Normal}(0,\sigma)\) distribution.

#### S.1.1 - Wasp size

After centering and standardizing to unit 1SD, tibia lengths \(z\_{m,i,j,k,o}\) with \(m\) the genetic mix of origin, \(i\) the experimental landscape of origin, \(j\) the source population (core, edge or stock) and \(k\) the individual (each individual measured twice, \(o\) denoting the observer) can be described by the following model:

\[
\begin{equation}
z\_{m,i,j,k,o} \sim \mathrm{Normal}(\mu\_{m,i,j,k},\sigma\_{r}), \\
\mu\_{m,i,j,k} = \beta\_{0} + \sum\_{n=1}^{N} \beta\_{n} \times x\_{n[m,i,j]} + \alpha\_{m} + \gamma\_{i} + \zeta\_{j} + \eta\_{k}, \\
\alpha\_{m} \sim \mathrm{Normal}(0,\sigma\_{\alpha}), \\
\gamma\_{i} \sim \mathrm{Normal}(0,\sigma\_{\gamma}), \\
\zeta\_{j} \sim \mathrm{Normal}(0,\sigma\_{\zeta}), \\
\eta\_{k} \sim \mathrm{Normal}(0,\sigma\_{\eta}). \\
\end{equation}
\]

In this model, the intercept \(\beta\_{0}\) denote the mean size in the stock populations, \(\beta\_{n}\) the other fixed-effect coefficients (here the connectivity × location-specific deviations from this starting size), and \(\alpha\), \(\gamma\), \(\zeta\), \(\eta\) random effects of genetic mix, experimental landscape, experimental population in landscape and individual identity, respectively. We used \(\mathrm{Normal}(0,1)\) priors for fixed effects (including the intercept), and \(\mathrm{Half-Normal}(0,1)\) priors for all standard deviations (including residual standard deviation \(\sigma\_{r}\)), following McElreath (2020).

#### S.1.2 - Short-term activity

For activity, we analyze the proportion of time spent active \(P\_{m,i,j,k}\), with again \(m\) denoting the mix of origin, \(i\) the landscape and \(j\) the population. \(k\) here corresponds to the replicate, as several independent sub-groups were tested per population of origin. For reasons outlined in the main text, we analyze replicate-level aggregate metrics, not individual-level traits. These proportions can be analyzed using a Beta model, which uses here the (mean, precision) parameterisation of the Beta distribution:

\[
\begin{equation}
P\_{m,i,j,k} \sim \mathrm{Beta}(p\_{m,i,j}, \phi), \\
\mathrm{logit}(p\_{m,i,j}) = \beta\_{0} + \sum\_{n=1}^{N} \beta\_{n} \times x\_{n[m,i,j]} + \alpha\_{m} + \gamma\_{i} + \zeta\_{j}, \\
\alpha\_{m} \sim \mathrm{Normal}(0,\sigma\_{\alpha}), \\
\gamma\_{i} \sim \mathrm{Normal}(0,\sigma\_{\gamma}), \\
\zeta\_{j} \sim \mathrm{Normal}(0,\sigma\_{\zeta}), \\
\end{equation}
\] Priors for fixed and random effects are as in **S.1.1**, except for the intercept \(\beta\_{0}\) which was set to \(\mathrm{Normal}(0,1.5)\); this follows suggestions from McElreath (2020) for parameters interpretable as the logit of a proportion. We set a \(\mathrm{Half-Normal}(0,1)\) prior on the inverse of the precision parameter (\(1/\phi\)), following suggestions made by Stan developers (https://github.com/stan-dev/stan/wiki/Prior-Choice-Recommendations).

#### S.1.3 - Effective dispersal models

Our measure of effective dispersal is the proportion of offspring \(p\_{m,i,j,k} = f\_{m,i,j,k}/F\_{m,i,j,k}\) found in the arrival patch, compared to the total number of offspring in a replicate \(k\) (arrival + departure patch, \(F\_{m,i,j,k}\)). This can be analyzed using binomial models:

\[
\begin{equation}
f\_{m,i,j,k} \sim \mathrm{Binomial}(F\_{m,i,j,k}, p\_{m,i,j,k}), \\
\mathrm{logit}(p\_{m,i,j,k}) = \beta\_{0} + \sum\_{n=1}^{N} \beta\_{n} \times x\_{n[m,i,j,k]} + \alpha\_{m} + \gamma\_{i} + \zeta\_{j}, \\
\alpha\_{m} \sim \mathrm{Normal}(0,\sigma\_{\alpha}), \\
\gamma\_{i} \sim \mathrm{Normal}(0,\sigma\_{\gamma}), \\
\zeta\_{j} \sim \mathrm{Normal}(0,\sigma\_{\zeta}). \\
\end{equation}
\] \(\beta\_{n}\) here include, in addition to the effects of connectivity × location, the effect of (standardised) total number of offspring and (in the model for the density-dependent experiment) effects of density and density × connectivity × location interactions. Priors for fixed and random effects are as in **S.1.1**, with the exception of the prior for the intercepts \(\beta\_{0}\). As this parameter denotes the logit of a proportion, we used \(\mathrm{Normal}(0,1.5)\) priors (McElreath, 2020).

#### S.1.4 -Fecundity models

The total number of hosts successfully parasitized by a wasp in 24h, \(F\), is best described by a zero-inflated negative binomial model, with \(p\) the probability of excess zeroes and \(\lambda\) the mean fecundity absent zero-inflation:

\[
\begin{equation}
F\_{m,i,j,k} \sim \mathrm{ZINegativeBinomial}(p\_{m,i,j,k},\lambda\_{m,i,j,k},\phi), \\
\log(\lambda\_{m,i,j,k}) = \beta\_{0[\lambda]} + \sum\_{n=1}^{N} \beta\_{n[\lambda]} \times x\_{n[m,i,j,k]} + \alpha\_{m} + \gamma\_{i} + \zeta\_{j}, \\
\mathrm{logit}(p\_{m,i,j,k}) = \beta\_{0[p]} + \sum\_{n=1}^{N} \beta\_{n[p]} \times x\_{n[m,i,j,k]} + \eta\_{m} + \nu\_{i} + \theta\_{j}. \\
\alpha\_{m} \sim \mathrm{Normal}(0,\sigma\_{\alpha}), \\
\gamma\_{i} \sim \mathrm{Normal}(0,\sigma\_{\gamma}), \\
\zeta\_{j} \sim \mathrm{Normal}(0,\sigma\_{\zeta}), \\
\eta\_{m} \sim \mathrm{Normal}(0,\sigma\_{\eta}), \\
\nu\_{i} \sim \mathrm{Normal}(0,\sigma\_{\nu}), \\
\theta\_{j} \sim \mathrm{Normal}(0,\sigma\_{\theta}). \\
\end{equation}
\] In the model for the density-dependent experiment, \(\beta\_{n}\) include, in addition to the effects of connectivity × location, effects of density and density × connectivity × location interactions.  
Priors for fixed and random effects are as in **S.1.1**, with three exceptions. First, the prior for the intercept of the negative binomial component (\(\beta\_{0[\lambda]}\)) was set to \(\mathrm{Normal}(3.8, 0.5)\). This is based on the fact wasps were offered about 90 eggs: this prior is centered on \(\log(45)\), and has only limited support for values > 90 when transformed back on the data scale. The prior for the intercept of the zero-inflation component (\(\beta\_{0[p]}\)) was set to \(\mathrm{Normal}(0,1.5)\); this follows suggestions from McElreath (2020) for parameters corresponding to the logit of a proportion. We use a narrower prior \(\mathrm{Half-Normal}(0,0.2)\) for the random effect SDs corresponding to genetic mix (\(\sigma\_{\alpha}\), \(\sigma\_{\nu}\)) here: the usual \(\mathrm{Half-Normal}(0,1)\) we use in all other cases induces a few divergences here, a not-surprising consequence of both the model being much more complex than the others and the limited number of mixes (3). The narrower prior helps convergence by regularizing; note that inferences are the same if we use the “standard” prior. Finally, we set a \(\mathrm{Half-Normal}(0,1)\) prior on the inverse of the shape parameter (\(1/\phi\)), again following suggestions made by Stan developers (https://github.com/stan-dev/stan/wiki/Prior-Choice-Recommendations).

### S.2 - Posterior pairwise comparisons between treatments

#### S.2.1 - Wasp size

**Figure S.2.1** - Posterior distributions of the pairwise between-context differences in predicted wasp size (tibia length). Black dots and segments are posterior means and 95% credible intervals; differences are given in SD units (1SD = 11.9 μm).

#### S.2.2 - Short-term activity

**Figure S.2.2** - Posterior distributions of the pairwise between-context differences in predicted probability of activity. Black dots and segments are posterior means and 95% credible intervals; comparisons are made on the logit scale.

#### S.2.3 - Effective dispersal

**Figure S.2.3** - Posterior distributions of the pairwise between-context differences in effective dispersal in the first (density-independent) experiment. Black dots and segments are posterior means and 95% credible intervals; comparisons are made on the logit scale.

**Figure S.2.4** - Posterior distributions of the pairwise between-context differences in effective dispersal in the second (density-dependent) experiment. Black dots and segments are posterior means and 95% credible intervals; comparisons are made on the log scale. Comparisons are made for each test density separately; for comparisons between low and high densities, see main text **Fig. 5C**.

#### S.2.4 - Fecundity

**Figure S.2.5** - Posterior distributions of the pairwise between-context differences in predicted reproductive success in the first (density-independent) experiment. Black dots and segments are posterior means and 95% credible intervals; comparisons are made on the log scale.

**Figure S.2.6** - Posterior distributions of the pairwise between-context differences in predicted reproductive success in the second (density-dependent) experiment. Black dots and segments are posterior means and 95% credible intervals; comparisons are made on the log scale. Comparisons are made for each test density separately; for comparisons between low and high densities, see main text **Fig. 6C**.

### S.3 - Effect of experimental connectivity level on demographic variability

In the main text, we mention that levels of environmental variability may influence the slope of the density-dispersal relationship, based on Rodrigues & Johnstone (2014), and that it *may* explain how core patches from landscapes with different connectivity differ in dispersal dynamics in unexpected ways.  
To validate this, we need to show that experimental treatment has an effect on environmental variability in core patches. In our case, this is equivalent to looking at demographic variability (because resources \_hosts\_ are fixed, so the amount of resource per individual is only driven by the number of individuals).  
We do have access to population size estimates for this experiment. These data are re-used from Dahirel et al. (2020), we invite the reader to look at the original source for details about the raw data characteristics. The processed dataset contains information about mean population size (proportion of hosts parasitised), SD and coefficient of variation CV for each experimental landscape. Because population sizes were estimated semi-automatically using four differently-biased computer macros that roughly compensate each other (again, see Dahirel et al., 2020), we have four values per landscape, one per macro/“observer.”

We analyse coefficients of variation using a lognormal mixed model:

\[
\begin{equation}
\mathrm{CV}\_{m,i,o} \sim \mathrm{LogNormal}(\mu\_{m,i,o}, \sigma\_{d}), \\
\mu\_{m,i,o} = \beta\_{0} + \beta\_1 \times \mathrm{TREATMENT}\_{[m,i]} + \alpha\_{m} + \gamma\_{o}, \\
\alpha\_{m} \sim \mathrm{Normal}(0, \sigma\_{\alpha}), \\
\gamma\_{o} \sim \mathrm{Normal}(0, \sigma\_{\gamma}), \\
\end{equation}
\]

where the \(CV\_{m,i,o}\) of replicate \(i\) from genetic mix \(m\) as seen by “observer”/macro \(o\) depends on its connectivity level and on random effects of mix and macro. We use \(\mathrm{Normal}(0,1)\) priors for fixed effects and \(\mathrm{Half-Normal}(0,1)\) for all \(\sigma\) (random effects and residuals).

We find that core patches were less demographically variable when connectivity was reduced, with a CV on average 0.68 [0.61, 0.74] times the one of reference patches (**Fig. S.3.1**).

**Figure S.3.1** - Posterior predicted demographic variability (Population size coefficient of variation) in core patches (\(x = 0\)) as a function of connectivity treatment. Black dots and segments are posterior means and 95% credible intervals, gray dots are observed values (averaged across “observer” macros).

### References

Dahirel, M., Bertin, A., Haond, M., Blin, A., Lombaert, E., Calcagno, V., … Vercken, E. (2020). Shifts from pulled to pushed range expansions caused by reduction of landscape connectivity. *bioRxiv, Version 4 Peer-Reviewed and Recommended by Peer Community in Evolutionary Biology*, 2020.05.13.092775. doi: 10.1101/2020.05.13.092775

McElreath, R. (2020). *Statistical rethinking: A bayesian course with examples in R and Stan* (2nd edition). Boca Raton, USA: Chapman; Hall/CRC.

Rodrigues, A. M. M., & Johnstone, R. A. (2014). Evolution of positive and negative density-dependent dispersal. *Proceedings of the Royal Society of London B: Biological Sciences*, *281*(1791), 20141226. doi: 10.1098/rspb.2014.1226
